# Abca4 inhibition in a cone-rich rodent leads to Stargardt Disease type 1-like retinal degeneration

**DOI:** 10.1101/2023.09.04.556201

**Authors:** Fabiana Sassone, Michel J. Roux, Dominique Ciocca, Paola Rossolillo, Marie-Christine Birling, Janet R. Sparrow, David Hicks

**Author notes:** Author for all correspondence : David Hicks, INCI-UPR3212-CNRS, 8 allée du général Rouvillois, 67000 Strasbourg, France.

## Abstract

Mutations in the gene *ABCA4* coding for photoreceptor-specific <u>A</u>TP-<u>b</u>inding <u>c</u>assette subfamily <u>A</u> member <u>4</u>, are responsible for the most common form of inherited macular degeneration known as Stargardt Disease type 1 (STGD1). STGD1 typically declares early in life and leads to severe visual handicap. *Abca4* gene deletion mouse models of STGD1 show increased accumulation of lipofuscin, a hallmark of the disease, but unlike the human disease show mostly no photoreceptor degeneration or functional decline (an albino *Abca4-/-* mouse exhibits photoreceptor degeneration although functional parameters were not studied). Reasoning that the small cone population of mice (<3%) might compromise more faithful modelling of human maculopathies, we performed subretinal injections of CRISPR/Cas9-Abca4 recombinant Adeno-Associated Virus constructs into young Fat Sand Rats (*Psammomys obesus*), a diurnal rodent containing >30% cones. Sanger sequencing of the CRISPR-targeted sequence showed clear edition of the *Abca4* gene. At 2 months post-injection, non-invasive fundus imaging showed widespread photoreceptor loss, confirmed by optical coherence tomography. Functional recording by scotopic and photopic single flash, and photopic flicker electroretinography, showed significant decline in photopic (cone) but not scotopic (rod) light responses. Post-mortem real-time PCR, immunohistochemistry and western blotting showed significant decrease of cone-specific (MW cone opsin) but not rod-specific (rhodopsin) markers. Transmission electron microscopy showed large numbers of lipid inclusions in treated but not control retinal pigmented epithelium. Finally, ultrahigh performance liquid chromatographic analysis of whole *P. obesus* eyes showed the presence of *all-trans* retinal-dimer, also seen in *Abca4*^-/-^ mice but not normal rod-rich mouse or rat eyes. In conclusion, this animal model of STGD1 more accurately reflects human STGD1 and should be valuable for characterizing pathogenic pathways and exploring treatment options.

## Introduction

Photoreceptor (PR) damage, malfunctioning and death occur in a large number of visual pathologies. Whether the underlying causes are genetic, chemical or environmental, progressive loss of these cells leads to visual handicap and incapacitating blindness. The death of cones is particularly devastating since these cells are responsible for high acuity and chromatic daylight vision [1]. Cones reach maximal density within the fovea, and this region is especially vulnerable in widespread diseases such as age-related macular degeneration and autosomal recessive maculopathy Stargardt Disease type 1 (STGD1). The latter is the most common inherited maculopathy, with carrier frequency of up to 1:50 in the general population. The causal gene has been identified as *ABCA4* encoding the PR-specific flippase <u>A</u>TP-<u>b</u>inding <u>c</u>assette subfamily <u>A</u> member <u>4</u> [2]. More than 1200 mutations have been identified, the majority of which are compound heterozygous, resulting in retinal degeneration of variable phenotype but in which early foveal damage and severe loss of central vision are commonly observed [3]. Most *ABCA4* mutations lead to a central-peripheral gradient of retinal damage, almost always with macular involvement [4, 5]. A commonly used classification recognizes three sub-types of STGD1: group 1 in which only macular degeneration is seen, with normal full-field rod and cone function; group 2 with macular degeneration and also pan-retinal cone malfunction, but mostly preserved rod function; and group 3 in which both cone and rod loss are widespread [5, 6]. The most severe forms are linked to null mutations, resulting in cone-rod dystrophy that appears early in life and evolves rapidly towards severe visual handicap. Abca4 is expressed by both rods and cones [7], and based on seminal studies using genetically modified mouse strains [8–10], biochemical analyses of retinal and pigmented epithelium (RPE) preparations [11] and rigorous *in vitro* experiments [12], a pathogenic mechanism has been proposed. Truncated or absent Abca4 impairs the visual cycle, with slowed clearance of *all-trans* retinal (*at*Ral) leading to formation of toxic bisretinoids within PR outer segments (OS). Daily phagocytic removal of PROS membrane debris by the adjacent RPE leads to accumulation and modification of these bisretinoids within the RPE, resulting in accelerated build-up of lipofuscin, a hallmark feature of STGD1 in humans and mouse models [3, 4, 8–10]. Lipofuscin exhibits noxious properties such as initiation of inflammatory cascades and photosensitization [13], and eventually toxicity reaches a threshold level at which RPE cells start to die; since RPE is critical for the maintenance of PR, these latter cells also die in their turn, forming a vicious pathogenic circle.

However, this hypothetical mechanism does not account for the particular phenotype seen in many STGD1 patients. As stated above, group 2 STGD1 patients exhibit pan-retinal loss of cone function while rod function remains relatively normal [5, 6]. Black and albino *Abca4* null mutant mice both display accumulations of bisretinoid lipofuscin [8–10] as seen in the human disease. Albino *Abca4-/-* mice also exhibit substantial PR loss at 8 months of age, the absence of melanin leading to increased photo-oxidative damage [13, 14]. Conversely black *Abca4^-/-^* mice show no structural or functional defects even in aged mice [8, 15]. Similarly, knock-in p.Leu541Pro/p.Ala1038Val mutant pigmented *Abca4* mice do not show any degenerative changes at 12 months [16], and black knock-in p.Asn965Ser mutant mice do not show retinal stress or PR loss at 2 months [17]. Finally, in severe forms of human STGD1 it has been reported that retinal abnormalities appear prior to visible changes in the RPE [18].

One potential drawback of relying on mouse models for maculopathies is the low numbers of cones in such nocturnal rodents: <1% in rats [19], <3% in mice [20]. We reasoned that animals containing high numbers of cones, more closely resembling the PR composition of human macula, would possibly constitute superior models for such diseases. In keeping with their temporal niche, diurnally active rodents possess high numbers of cone PR, ∼30-40% for those murid species studied [21], in which rod and cone cell bodies are separated into distinct sublayers in the outer nuclear layer (ONL), facilitating their analysis [21, 22]. Furthermore, their visual response characteristics are more similar to those of humans compared to rats or mice, with a robust photopic response [23, 24]. We show here that knockout of *Abca4* transcription in a cone-rich rodent leads to widespread and rapid PR degeneration with prominent structural and functional decline in cones, more closely echoing the STGD1 phenotype.

## Results

### Confirmation of *Psammomys* Abca4 genomic sequence

The region containing the target by CRISPR/Cas9 was confirmed by Sanger sequencing and confirmed significant similarity with the *Acomys cahirinus* (479/571, 84%) [25] and *Rattus norvegicus* (412/508, 81%) [26] genomic sequences.

### Verification of viral *abca4* CRISPR/Cas9-mediated edition in rod and cone PR

We used sub-retinal injection of recombinant adeno-associated viruses (rAAV2/8) to deliver *sp*Cas9 and *Abca4* gRNAs in order to create an *abca4* knockout in the Fat Sand Rat *Psammomys obesus*. We initially characterized endogenous Abca4 protein expression in postnatal and adult animals, and observed that expression levels were relatively low, compared to another cone-specific marker cone transducin (GαT2), prior to PN20 (Figure 1A-A’’’, B-B’’’), and in terms of their mRNA and protein expression profiles (Figures 1C, D). This provided a time window of ∼5d to knock down the gene between eye opening (∼PN13) and prior to high levels of endogenous protein. We verified that rAAV2/8 serotype efficiently and selectively transfected both rod and cone PR by sub-retinal injection of eGFP-tagged rAAV2/8 into PN15 pups: we observed widespread transfection of both rods and cones but no other retinal cells in other cell layers (Figure 1E-E’’). For *Abca4* editing, due to the limited cargo capacity of AAV vectors it was necessary to co-inject two viruses, one containing sgRNAs targeting *Psammomys obesus Abca4* and an eGFP expression cassette, and the second containing an HA-tagged Cas9, into the sub-retinal space of young (PN14-17) animals (Figure 1F, G). We chose to use a double guide approach, one guide (gR58) directing a double strand break at the 5’ of exon 5, the other guide (gR80) directing a double strand break just after the 3’ end of exon 5. Ideally, if both guide RNAs were efficient, the whole exon 5 would be deleted, leading to a frame shift (premature STOP codon) and knock- out of the gene. The close proximity of the splice donor or splice acceptor of exon 5 of both guides ensured a KO allele even if only one guide was efficiently generating a double strand break.

**Figure 1.**
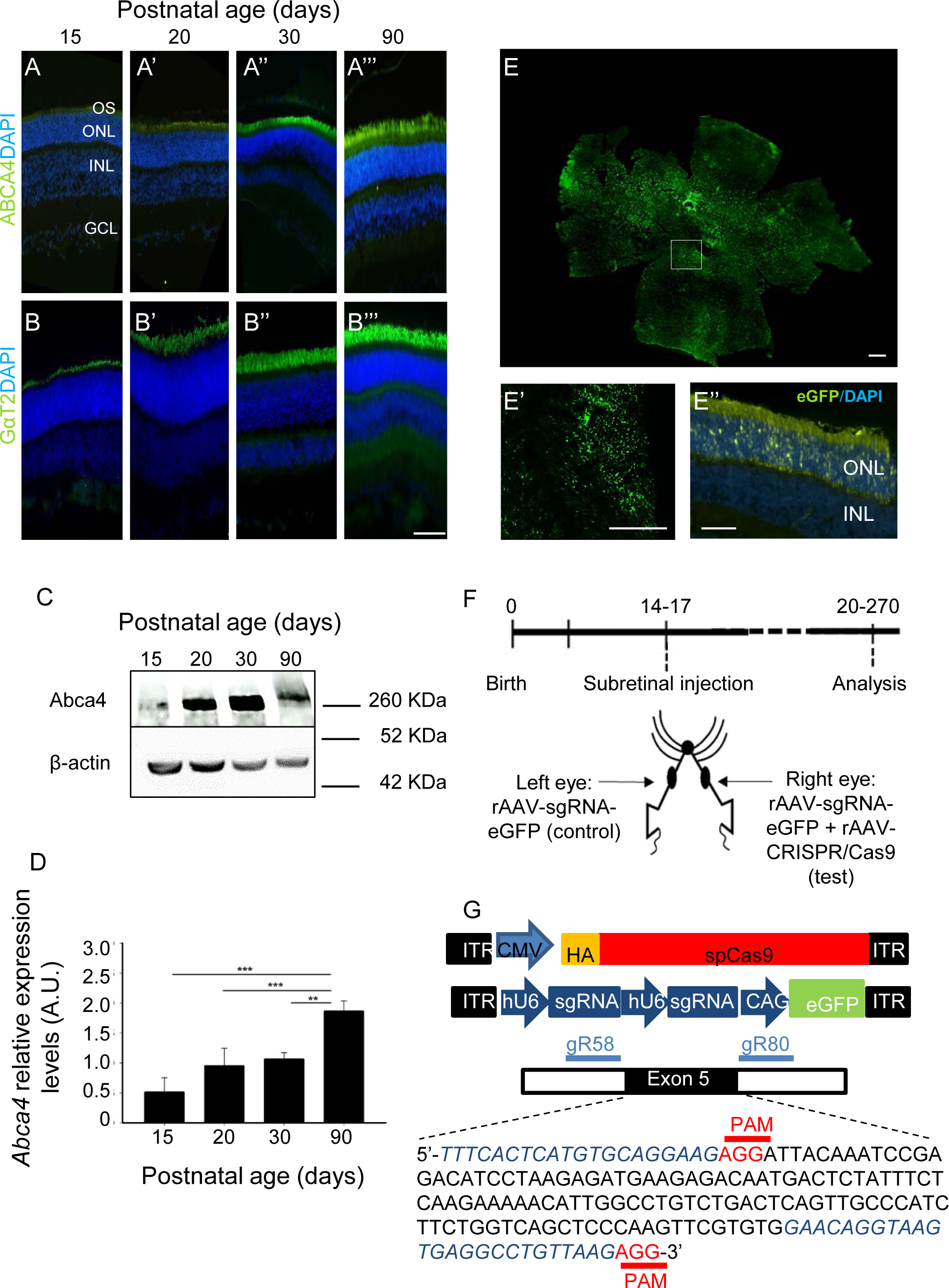
Abca4 expression in Psammomys obesus retinal development and subretinal injection of rAAV-CRISPR-Cas9. (A-B) Immunostaining of abca4 (A-A’’’, green) and cone transducin (GαT2) (B-B’’’, green) in retinal tissues collected from *Psammomys obesus* aged postnatal day P15, P20, P30 and P90. At all ages, whereas GαT2 labelling is already strong, abca4 is only faintly visible at P15 and 20, becoming more intense by P30. Abbreviations: DAPI, 4′,6-diamidino-2-phenylindole; GCL ganglion cell layer; INL inner nuclear layer; ONL outer nuclear layer; OS outer segments. Scale bar in B’’’ = 40µm for all panels. (C) Western blotting of abca4 in P15-90 *P. obesus* retina. Compared to the housekeeping gene β-actin, abca4 immunoreactivity increased with postnatal age. (D) Quantification of abca4 immunoreactivity in P15-90 *P. obesus*. Densitometric scanning of abca4 immunostained bands, normalized to β-actin, showed significant increases in expression levels with increasing age, n of 3 independent experiments. (E) Representative retinal whole-mounts (top panel, insert shown as white box at higher magnification in E’) and sections (E’’) of a treated animal 7d post-injection. EGFP expression (green) indicates transcription by the rAAV-sgRNA vector in the PR layer. Abbreviations: IS, inner segments; ONL, outer nuclear layer. Scale bars = 100µm for panels E and E’, 40µm for panel E’’. (F) Timeline for the Abca4 knockdown experiments. *Psammomys obesus* received subretinal co-delivery of 1.4 x 1^10^ vector genomes (vg) rAAV-Cas9 and 1.4 x 1^10^ vg rAAV-sgRNAs-EGFP per eye at P14-P17. Same doses of rAAV-sgRNAs-EGFP were injected in fellow eyes as controls. (G) Schematic representation of the AAV vectors delivering SpCas9 and SgRNAs. Abca4 locus showing the location of the sgRNA targets (gR58, gR80). The targeted genomic site is Exon 5 in black. Protospacer adjacent motif (PAM) sequence is marked in red.

Next, in order to confirm CRISPR/Cas9 gene editing, retinas were collected 4 weeks post-injection and processed by fluorescent-activated cell sorting (FACS) to isolate eGFP-tagged control (gRNA only) and test CRISPR/Cas9 PR (Figure 2A). We estimated ∼25% total PR were transduced by counting eGFP-positive cells in fluorescence-activated cell sorted (FACS)-retina (Figure 2A). Agarose gel analysis showed the presence of a ∼250bp band shift in test cells only (Figure 2B), and Sanger sequence analysis of this band confirmed a deletion in *Abca4* gene, with almost no WT sequence remaining (Figure 2C). In further tests of the efficiency of *Abca4* knockdown, immunohistochemical staining of retinas at 2 months post-injection showed that compared to sham-injected eyes, those receiving CRISPR/Cas9 rAAV exhibited qualitatively reduced Abca4 label (Figure 2D-D’’, E-E’’); Cas9 immunostaining was restricted to transduced PR in animals injected with active viral probe (Supp. Figure 1). Finally, the evaluation of *Abca4* mRNA levels by PCR in control and test eyes showed significant decreases only in test eyes (Figure 2F); and western blots against Abca4 showed reduced band intensity in test but not control retinas (Figure 2G).

**Figure 2.**
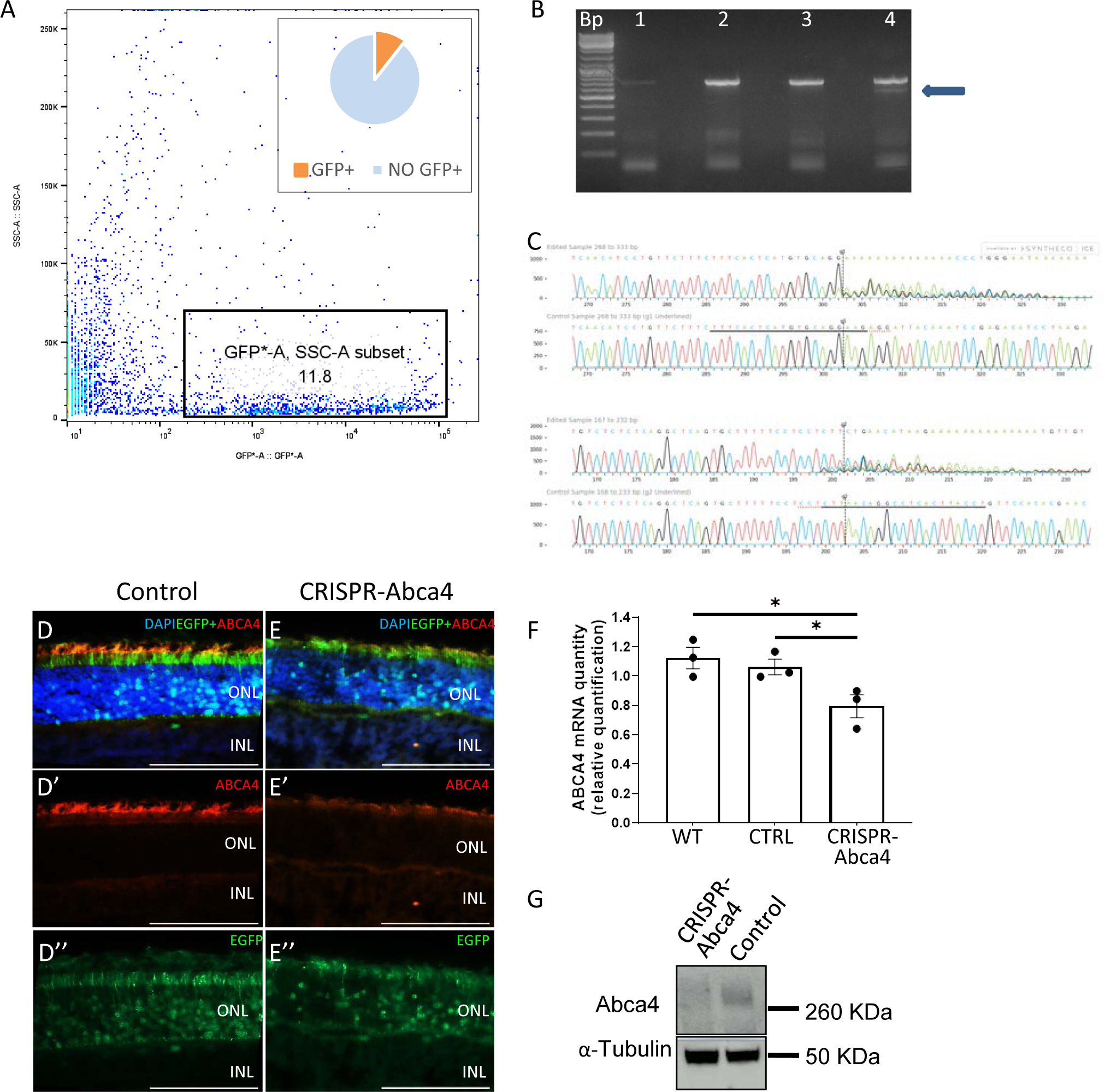
Abca4 knockdown in Psammomys obesus photoreceptors by AAV-CRISPR-Cas9 gene editing. (A) Representative FACS plots of dissociated cells from retinas receiving CRISPR-Abac4 vectors. EGFP-positive population represents 11.8% of total retinal cells. (B) Electrophoresis of *Abca4* cDNA following PCR amplification (∼700bp around exon 5): Bp, base pair ladder; 1, negative control (water only), no amplification; 2, positive control, *P. obesus* whole retina, expected band ∼700bp; 3, *P. obesus* retina transfected with rAAV gRNA/eGFP only, FACS-sorted cells show single band ∼700bp; 4, *P. obesus* retina transfected with rAAV gRNA/CRISPR/Cas9, FACS-sorted cells show two amplicons, one with edited *Abca4* (∼500bp) (blue arrow) and the other with wild type *Abca4* (∼700bp). (C, C’) Sanger sequencing chromatograms from forward primer (C) and reverse primer (C’). In each condition, we observed flattening of peaks and loss of correlation with DNA sequence in FACS-sorted cells starting from nucleotide ∼200-300 up to nucleotide ∼300-400, ie. the exon 5 region targetted by sgRNA-CRISPR/Cas9, compared to FACS-sorted control cells with a normal chromatogram profile, shown in the lower traces. (D, E) Immunostaining of Abca4 in retinal tissues collected from *Psammomys obesus* retina, rAAV-sgRNA-eGFP (control) (D- D’’’) and rAAV-sgRNA/eGFP+rAAV/ CRISPR/Cas9 (treated) (E-E’’’), 8 weeks post injection. Panels D, E: merged images of DAPI staining, abca4 immunostaining and eGFP. Panels D’, E’, substantial reduction of Abca4 immunostaining was observed in the CRISPR/Abca4- treated retina compared to control. Panels D’’, E’’, eGFP signal in transduced PR. Scale bar in E’’ = 80µm for all panels. Abbreviations: INL inner nuclear layer; ONL outer nuclear layer. (G) Quantitative real-time RT–PCR analysis of *abca4* mRNA expression in *Psammomys obesus* retinas 8 weeks post-injection rAAV-sgRNA/eGFP+rAAV-CRISPR/Cas9 (treated) (n=3), rAAV-sgRNA/eGFP control retinas (n=4), and unoperated retinas (n=3). The mRNA expression values were determined after normalization to internal control GAPDH and RPLP0 mRNA levels. CRISPR-treated vs. control p=0.024 CRISPR-treated vs. WT p=0.011, one way ANOVA. Data represented as mean ± SEM. *P < 0.05. (G) Representative immunoblot of Abca4 (260kDa) in *Psammomys obesus* rAAV-sgRNA/eGFP+rAAV/ CRISPR/Cas9 (treated) retinal lysate, and unoperated retinal lysate at 270 days post injection. α tubulin (50kDa) as loading standard.

### Abca4 loss of function leads to decline in cone but less rod visual responses

Scotopic single flash ERG recordings of CRISPR/Cas9-treated (test) and sgRNA/eGFP virus-injected (control) eyes 2 months post-injection (∼PN75) showed typical stimulus intensity/response amplitude relationships. Weak flashes (pure rod responses) produced stable linear “a” wave traces, increasing at higher flash intensities (mixed rod/cone responses) (Figure 3A). There was a significant decline in “a” wave amplitudes at higher stimulus intensities in CRISPR/Cas9 treated but not control-injected controls (Figure 3A). There was a similar decline in scotopic “b” wave amplitudes, particularly at higher intensities (Figure 3B). Photopic (pure cone) responses also showed significant decreases in amplitude, for both “a” and “b” waves (Figures 3C, D). Representative single scotopic and photopic traces are shown in Supp. Figures 2 and 3 respectively. We also performed photopic flicker ERG recordings, and there too we saw significant reductions in peak amplitudes only in CRISPR/Cas9-treated animals (Figure 3E; Supp. Figure 4). No significant differences were observed when comparing rAAV-sgRNAs/eGFP-injected (control) and non-operated eyes (Supp. Figure 5). Similar data were obtained for animals after 90 and 210d post-injection, which showed that cone “a” and “b” wave, and also rod “b” wave amplitudes, continued to decline over time (Supp. Figure 6).

**Figure 3.**
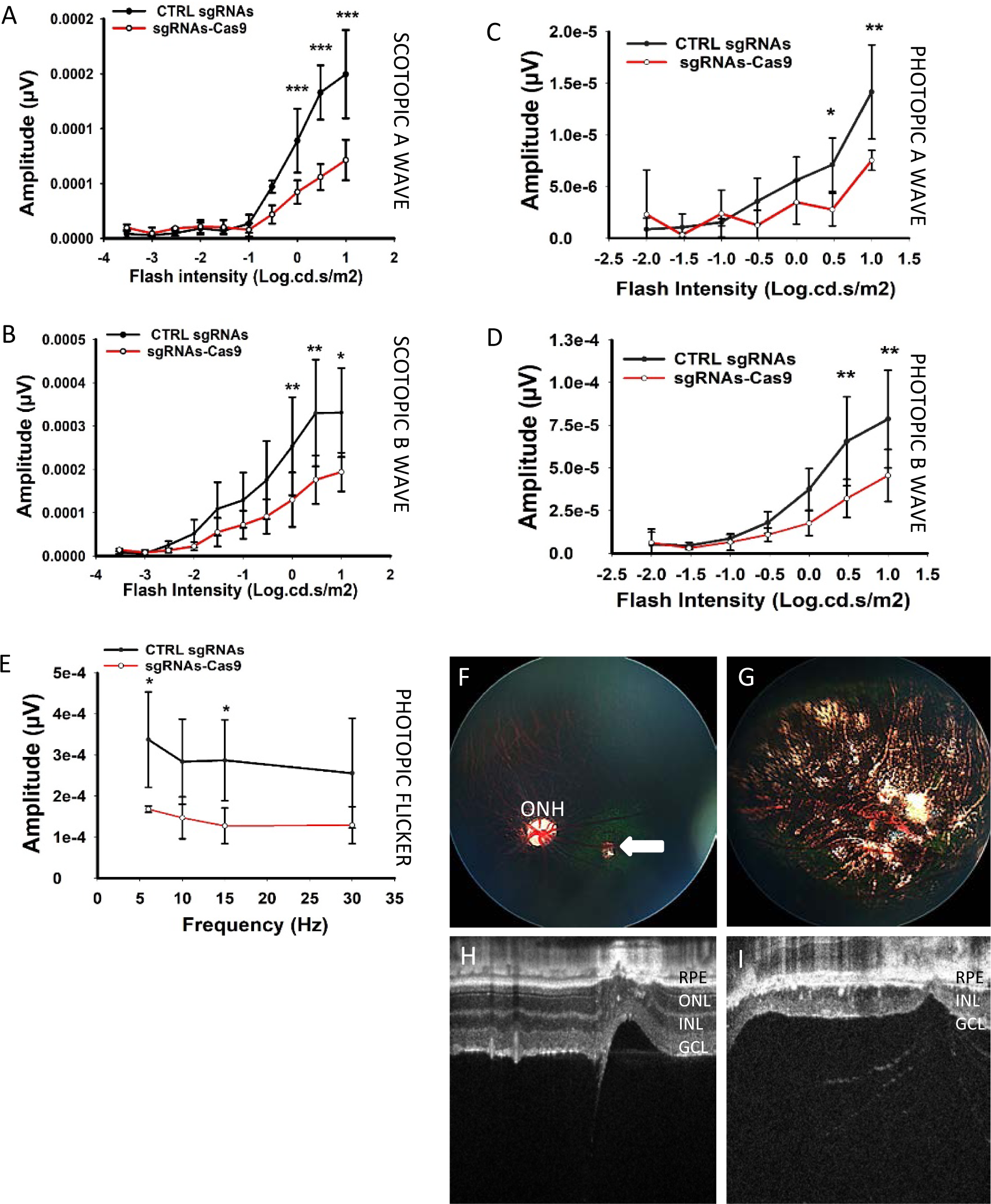
Morphological and functional changes in *Psammomys obesus* retina following Abca4 knockdown. (A-D) Mean ERG amplitudes recorded across a range of light flashes under scotopic or photopic conditions. Statistically significant lower amplitudes of scotopic a- and b-wave (A, B) (n=6), and photopic a- and b- wave (C, D) (n=3) were obtained in response to increasing intensities of flash stimuli. Values from CRISPR/Cas9-treated eyes (left eye, red trace) were compared to rAAV-sgRNAs/eGFP control eyes (right eye, black trace). There were significant differences between treated and control eyes at light flash intensities of ∼0.5-1 log.cd .s/m^2^ for scotopic conditions, intensities at which a mixed rod-cone response is present. Significant differences between treated and control eyes in photopic conditions were seen for light flash intensities 0.5-1 log.cd .s/m^2^. Data represented as mean ± SEM. *P < 0.05; **P < 0.01; ***P < 0.001. Significance values between CRISPR/Cas9-treated and control sgRNA/eGFP eyes was calculated using two way ANOVA. (E) Photopic light-adapted flicker recordings were made from 5 to 30 Hz, and also showed reduced signals in CRISPR/Cas9-treated eyes. Statistical analysis performed as for single flash experiments. (F-I) Fundus imaging of CRISPR/Cas9-treated and sgRNAs/eGFP control eyes at 8 weeks post-injection. Control eyes showed a small lesion corresponding to the injection site (F, white arrow), but otherwise showed smooth uniform bluish retina with blood vessels emanating from the optic nerve head (ONH); G, Optical Coherence Tomography (OCT) imaging of control retinas showed the normal laminated retinal structure, with a brightly reflective retinal pigmented epithelium (RPE) and dark cell layers (INL, inner nuclear layer; and ONL, outer nuclear layer) with the GCL (ganglion cell layer) separated by a wide plexiform layer; H, fundus imaging of CRISPR-CAS9-treated eyes showed extensive loss of retina with pigmented patches and visible blood vessels; (I) OCT imaging of CRISPR/Cas9-treated eyes showed a complete disappearance of the ONL, while vestigial inner retina was still present.

### Abca4 loss of function causes widespread photoreceptor degeneration

Fundus imaging of animals at 2 months after injection with control rAAV-sgRNA/eGFP showed a very small lesion corresponding to highly localized damage at the site of injection (arrow, Figure 3F). Otherwise the retinal aspect was uniform and normal, with major blood vessels faintly visible (Figure 3F). On the other hand, animals injected with active CRISPR/Cas9 probes showed widespread degeneration after 2 months, with extended depigmented lesion areas and blood vessels appearing under the damaged tissue (Figure 3G). Patches of fluorescence were visible at the lesion borders, indicating continued viral eGFP expression. OCT sections of control-injected animals showed the normal laminated structure of retina, with clearly visible cellular and synaptic layers (Figure 3H); however, CRISPR/Cas9-treated animals displayed severely disrupted tissue, the neural retina either missing entirely or just residual inner retina lacking the outer nuclear layer (ONL) (Figure 3I). Additional examples of fundus and OCT imaging of CRISPR/Cas9-injected eyes are shown in Supp. Figures 7-9. Longitudinal follow-up of animals 15, 30 and 60d after subretinal injection showed that eyes receiving control virus exhibited a small lesion due to the injection injury, which did not evolve over time; on the other hand, eyes receiving rAAV-CRISPR/Cas9 showed progressive retinal changes between 30 and 60d, with enlargement of lesions (Supp. Figure 9).

Immunohistochemical staining using a rod-specific antibody marker (rhodopsin, RHO) of 2 month post-injection control-injected and CRISPR/Cas9-injected retinas (the latter at areas close to the lesion borders, with continued eGFP expression in many cells) showed that rhodopsin label intensity was approximately equal in both cases (Figures 4A-A’’’, B-B’’’: compare Figures 4A’’’ and 4B’’’). This similarity was confirmed by quantitative PCR analysis of RHO mRNA in control and test retinas (Figure 4C). In contrast, use of a cone-specific antibody marker (mid wavelength-sensitive cone opsin, OPN1MW) to label 2 month post-injection control-injected and CRISPR/Cas9-injected retinas (Figures 4D-D’’’, E-E’’’) showed reduced OPN1MW signal in CRISPR/Cas9 compared to control eyes (compare Figures 4D’’’ and E’’’). This decrease was confirmed by quantitative PCR analysis of OPN1MW mRNA in control and test retinas (Figure 4F). Ultrastructural observation of the OS/RPE interface by transmission electron microscopy showed that control animals displayed conventional subcellular organization, with well-aligned rod and cone OS abutting the RPE (Figures 4G-I). RPE contained numerous oblong melanosomes, together with occasional phagosomes (ingested OS membranes), along their apical surface (Figures 4G, H). The subcellular aspect of the different cell regions, RPE, OS and inner segments (IS) showed good integrity and well-preserved organelles (Figures 4G-I). On the contrary, examination of CRISPR/Cas9 treated eyes areas close to lesion borders revealed RPE congested with numerous circular lipid inclusions (Figure 4J, K). Furthermore, many IS and OS displayed disrupted cytoplasm and loss of cellular integrity (red arrows, Figures 4J, L).

**Figure 4.**
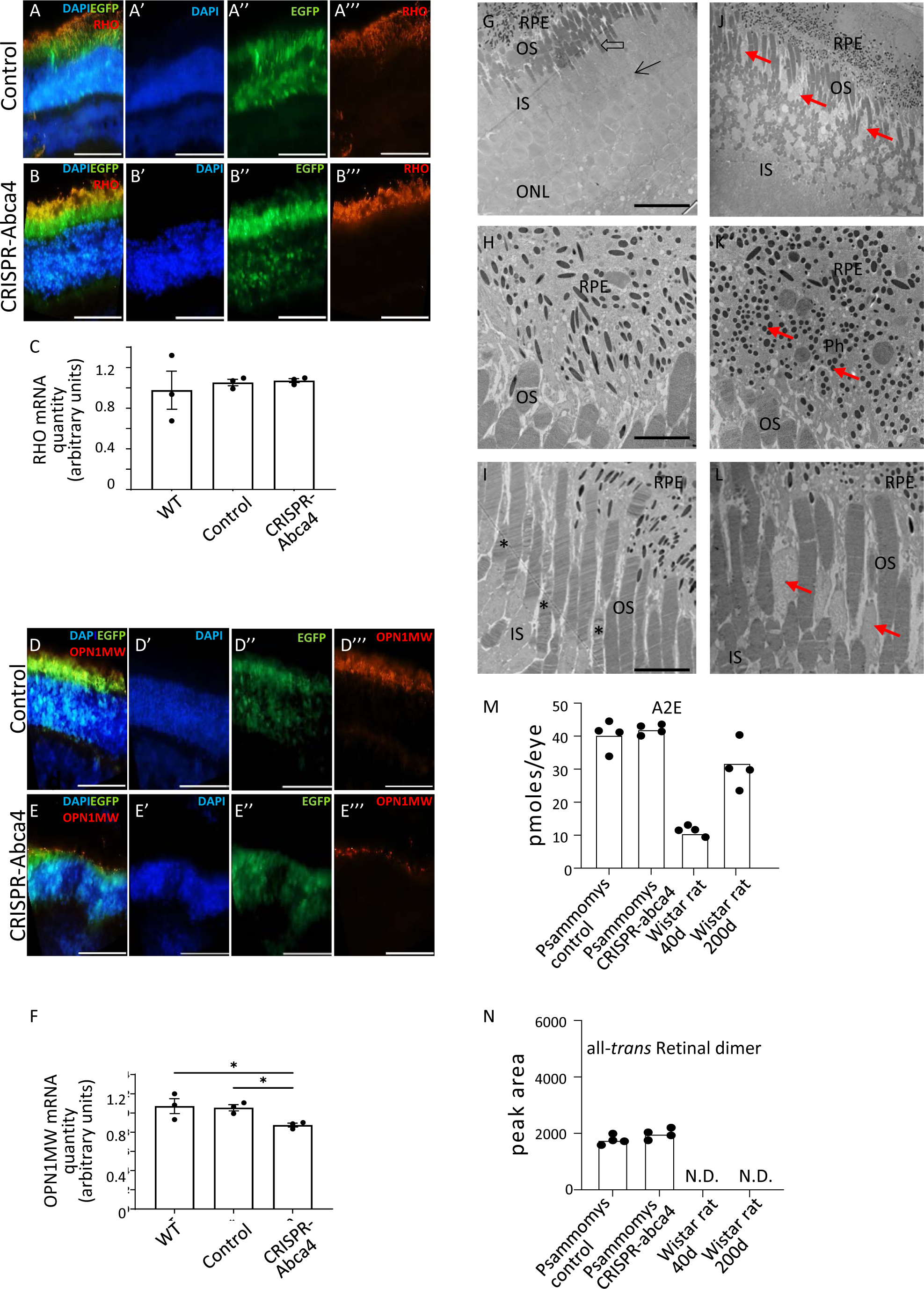
Rod, cone and RPE changes following Abca4 knockdown in *Psammomys obesus*. (A, B) Immunostaining of rhodopsin (RHO) in retinal tissues collected from *Psammomys obesus,* rAAV-sgRNA/eGFP (control) (A-A’’’) and rAAV-sgRNA/eGFP+rAAV/CRISPR/Cas9 (treated) (B-B’’’) at 8 weeks post injection. Sections were made close to lesion edges, where residual retina was still present. A, B: merged images of DAPI staining (blue), eGFP expression (green) and RHO immunostaining (red). DAPI (A’, B’), eGFP (A44, B’’) and RHO (A444, B’’’) alone do not show major differences between control and treated tissue. Scale bars = 40µm. (C) Quantitative real-time PCR analysis of *rho* mRNA expression in *Psammomys obesus* retinas 8 weeks post-injection. rAAV-sgRNA/eGFP+rAAV/ CRISPR/Cas9 (treated) (n=4), rAAV-sgRNA-EGFP control retinas (n=4), wild type retinas (n=3). The mRNA expression values were determined after normalization to internal control GAPDH and RPLP0 mRNA levels. There is no statistically significant difference between CRISPR-treated and either eGFP control or wild type retina (P = 0.915). One way ANOVA. (D, E) Immunostaining of OPN1MW in retinal tissues collected from *Psammomys obesus*, rAAV-sgRNA/eGFP (control) (D-D’’’) and rAAV-sgRNA-eGFP+AAV-CRISPR/Cas9 (treated) (E-E’’’) at 8 weeks post injection. D, E: merged images of DAPI staining (blue), eGFP expression (green) and OPN1MW immunostaining (red); whereas DAPI (D’, E’) and eGFP (D’’-E’’) alone do not differ between the two samples, there is a large decline in OPN1MW immunostaining in treated compared to control retinas (D’’’, E’’’). Scale bars = 40µm. (**F**) Quantitative real-time PCR analysis of *opn1mw* mRNA expression in *Psammomys obesus* retinas 8 weeks post-injection. rAAV-sgRNA/eGFP+rAAV-CRISPR/Cas9 (treated) (n=4), rAAV-sgRNA/eGFP control retinas (n=4), wild type retinas (n=3). The mRNA expression values were determined after normalization to internal control GAPDH and RPLP0 mRNA levels. CRISPR-treated vs. control p=0.036, CRISPR-treated vs. WT p=0.027. One way ANOVA. Data represented as mean ± SEM. *P < 0.05. (G-L) Transmission electron microscopy of the RPE/PR interface in rAAV-sgRNA/eGFP control (G-I) and rAAV- CRISPR/Cas9-treated eyes (J-L) at 27 weeks post-injection. G, low magnification images of the entire area showed normal ultrastructural features: RPE contained elongated melanosomes closely adjacent to PR OS, the rod OS appearing cylindrical, more darkly staining and closer to the RPE surface, whereas cone OS were more deeply embedded and lighter stained (open arrow). The inner segments (IS) were demarcated from the underlying cell bodies in the ONL by the outer limiting membrane (arrow). H, higher power image of RPE/OS interface, again showing numerous elongated melanosomes within the RPE, as well as occasional phagosomes. The rod OS show normal stacked discs. I, higher power image of OS/IS junction, again the OS are intact and neatly aligned, the cones OS showing a tapered aspect (*). J, low power images of RPE/PR region in CRISPR-CAS9 treated eyes. The RPE contains numerous circular inclusions packed into the apical cytoplasm, and many holes and damaged cells are seen in the OS layer, particularly at the level of the cone OS (red arrows); K, higher power images of RPE/OS interface, highlighting the abundant small round deposits packing the RPE (red arrows), as well as several phagosomes. These deposits form a dense layer between the deeper RPE containing melanosomes and the abutting OS; L, higher power images of the OS/IS region, showing the scattered free cytoplasm from degenerated cells and spaces between OS. Again, the rod OS appear less damaged than the cone OS, which are no longer recognisable (red arrows). Scale bar = 10µm (G, J), 2µm (H, I, K, L). (M) UPLC quantification of the bisretinoid A2E from whole eyes of 2 month post-injection *Psammomys obesus* (n=4) and albino Wistar rats *Rattus norvegicus* at 40 and 200 days (n=4 each). Data represented as mean ± SEM. High levels of A2E are detected in *P. obesus* compared to Wistar rats, in both control and CRISPR/Cas9- treated eyes. (N) UPLC quantification of all-trans Retinal dimer (atRal-di) from whole eyes of 2 month post-injection *Psammomys obesus* (n=4) and albino Wistar rats *Rattus norvegicus* at 40 and 200 days (n=4 each). *P. obesus* contain significant amounts of this molecule, whereas it is undetectable in Wistar rats. Data represented as mean ± SEM.

Ultrahigh performance liquid chromatography (UPLC) of extracts prepared from whole albino Wistar rat eyes, or control- or CRISPR/Cas9-injected *Psammomys* eyes, showed large differences in two key bisretinoids: A2E and *at*Ral-dimer. Although there was no difference between control and test *Psammomys* eyes (2 month post-injection, ∼75d old animals), with both having ∼41 pmoles/eye A2E, they contained significantly more A2E than either young (∼40d, ∼11 pmoles) or old (∼200d, ∼32 pmoles) rat eyes (Figure 4M). Additionally, while both groups of *Psammomys* eyes contained high amounts of *at*Ral-dimer, this bisretinoid was below detection limits in both groups of rat eyes (Figure 4N).

## Discussion

STGD1 is a devastating inherited blinding disease, the more severe forms declaring in early childhood and progressing rapidly to extreme visual handicap. Although knockout mouse models have existed for many years [8–10] (and more recently knock-in mutant *Abca4* transgenic mice: 16, 17), and exhibit some features of the disease (notably lipofuscin accumulation in the RPE, and age-related PR loss in the albino *Abca4*^-/-^ strain), but these animals have few cones. In the present study, knockdown of *Abca4* in the cone-rich rodent *Psammomys obesus* led to rapid and profound degenerative changes in the PR and RPE, and preferentially affected cone function and survival after 2 months. This is despite the transfection of only ∼25% total PR as opposed to complete germline ablation in the mouse studies. A major difference between these two types of models is the percentage of cones, <3% in mice [20] compared to ∼33% in *Psammomys* [22]. We speculate that in such diurnal rodents the retina basically constitutes one whole “macula-like” tissue in terms of cone density. This could provide an enormous advantage for better understanding the detailed etiology of macular diseases, not limited to STGD1 but possibly also relevant for pathologies like age-related macular degeneration.

Based on the knockout mouse models, clinical observations and extensive biochemical analyses, a hypothetical scenario for PR death in STGD1 has been proposed: under normal conditions, following light activation of rhodopsin, lipophilic *at*Ral separates from the opsin protein and remains within the luminal face of disc membranes. There it reacts with phosphatidyl-ethanolamine (PE) to form N-retinylidene-PE (NRPE), which is transported rapidly into the OS cytosol by the flippase ABCA4 localized in the disc periphery [12]. Once in the PR cytosol, *at*Ral is reduced by retinol dehydrogenase to *all-trans*-retinol, whereupon it is transported across the sub-retinal space to the adjacent RPE to complete re-isomerization to *11-cis* retinal *via* the visual cycle [27]. In STGD1, mutations in *ABCA4* lead to reduced or absent transport of NRPE; excess *at*Ral remaining within the disc lumen further reacts with PE to form a complex mixture of irreversible bisretinoid adducts. Bisretinoids are toxic bi-products of the visual cycle, including but not limited to A2E, exhibiting both photo-reactive and detergent properties [13]. Since PR undergo continuous membrane turnover involving apical displacement of discs and their internalization by the adjacent RPE [24], this latter cell type accumulates these toxic bisretinoids which cannot be degraded. They incorporate into lipofuscin deposits in RPE, and aggregations of bisretinoid lipofuscin in STGD1 are visible as conspicuous autofluorescent flecks by ophthalmological examination [28]. Eventually lipofuscin concentrations reach a toxic threshold causing the RPE to die, the metabolic maintenance of PR integrity is lost and the PR then die in turn. Detailed clinical studies of cohorts of STGD1 patients have established that enhanced lipofuscin accumulation appears as the earliest disease manifestation, and that rod and cone sensitivity (death) occur simultaneously [29].

Still, such a mechanism does not account for the preferential cone malfunction and loss seen in group 2 STGD1, and we propose a modified pathogenic sequence to take this into account. We recently demonstrated that cones exhibit several distinct molecular differences compared to rods: cones contain significantly more *Abca4* mRNA and Abca4 protein; they are highly deficient in very long chain fatty acids in general, and the neuroprotective poly-unsaturated fatty acid docosahexaenoic acid (DHA, 22:n:6) in particular; they contain far less PE; and show reduced amounts of A2E and five-fold greater amounts of *at*Ral-dimer [30]. Additionally, phagocytic turnover of cones by the RPE is much slower than rods [21]. We hypothesize that these compositional and functional differences render cones more vulnerable to cellular stress such as oxidative damage, and that disease mutations in STGD1 would lead to preferential cone death through direct poisoning, in addition to the indirect route described above. We speculate that because of the compositional differences, Abca4 failure in cones leads to the formation of bisretinoid species - such as *at*Ral-dimer - distinct from those detected in rods. Published data indicate *at*Ral-dimer levels are low in normal rod-dominated retinas but 3-4 fold higher in *Abca4^-/-^*mice compared to wild-type mice, and *at*Ral-dimer generates increased levels of singlet oxygen compared to A2E [31]. This indicates that reduced clearance of *at*Ral via defective Abca4 function leads to dimerization. Our data suggest they also constitute an important component of cone bisretinoids, possibly due to the low PE content pushing the equilibrium away from NRPE formation towards *at*Ral-dimer condensation. Increased levels of this toxic compound could explain the heightened vulnerability of cones seen in STGD1 and the present study. Intriguingly, a double knockout *Nrl/Abca4* mouse strain (all cone background) showed distinct differences in bisretinoids compared to the single *Abca4* knockout strain in wild-type mice (rod background): Abca4-deficient cones simultaneously generated more A2E than rods but were less able to clear it, suggesting that primary cone toxicity may occur [32]. A possible pathogenic pathway could come from Abca4-induced perturbations in retinoid recycling leading to membranes “congested” with *at*Ral-dimer (among other bisretinoids), with as a consequence reduced availability of *11-cis* retinal to reinsert into the OS. Different mouse models in which genes involved in the visual cycle were deleted have shown that reduced availability of *11-cis* retinal impacts cone survival and function more severely than rods [33–35]. Tying together the phenotype seen in *Psammomys* at 2 months after Abca4 knockdown (extensive areas of total PR loss, residual areas still containing transfected rods but not cones) and clinical features of type 2 STGD1, we suggest that *Abca4* mutations are highly toxic for cones but less for rods. Although we emphasize the preponderant effect of Abca4 knockdown on cones in the present study, rods do also die in STGD1, and we underline that rods also degenerate in *Psammomys*, but later than cones since scotopic ERGs declined between 3 and 7 months post-injection (Suppl. Figure 6).

In the present study, the method used (intra-ocular injection of virally-delivered CRISPR/Cas9) leads to a mosaic of untransduced (normal) and transduced (homozygous or heterozygous KO) rod and cone PR. This provides an experimental setting to test any differential effects of *Abca4* loss upon rod and cone survival, and permits use of the twin eye as an internal control. Even though our estimates show a maximum of ∼25% PR were transduced, *Abca4* knockdown results in widespread cell loss, already detectable after two months. Although we could not totally rule out a possible off-target effect, since the fully annotated genome of *Psammomys obesus* is not yet available, we believe this is highly unlikely for several reasons: 1. we based our gRNA design on the rat genome, which is highly homologous to that of *Psammomys* (Sanger sequencing of the *Abca4* coding region = 94% identity, 81% including intronic sequences), and the CRISPOR programme failed to identify any sequences recognized by the selected gRNA within the rat genome; 2. PCR analysis of *Abca4* mRNA levels, and western blotting and immunohistochemistry of Abca4 protein, showed clear reductions in treated eyes; 3. the total number of animals showing retinal degeneration (n=16) makes it highly unlikely off-target effects happened in every case. The data clearly suggest cones are more affected than rods at these early times, in terms of both structure (reduced expression of cone- but not rod-specific genes) and function (decreased photopic and flicker ERGs). We further postulate that such direct cone poisoning may have important implications for clinical strategies, since it would suggest that therapies involving stem cell replacement of RPE [36] may help rescue rods which die secondarily to RPE cells, but be of less value for cones which die by direct poisoning.

In conclusion, cone-rich rodents offer a unique scenario to explore molecular and cellular changes occurring in human maculopathies like STGD1, and should provide a valuable means to further explore genotype-phenotype relations, and to appraise potential therapeutic strategies.

## Materials and Methods

### Animals

*Psammomys obesus* were originally imported from the laboratory of Dr. Kronfeld-Schor, Tel Aviv, Israel, and have since been maintained as a viable breeding colony in the Chronobiotron CNRS UMR3415, in controlled ambient illumination on a 12 h light/12 h dark cycle, in environmentally enriched cages and with access to low caloric-density food which does not induce diabetes [37] (Altromin International, Germany) and water *ad libitum*. Animal use in the present project was authorized by local ethics committees (CREMEAS and Com’Eth) and the French Ministry of Research (APAFIS 8472-2016121318254073 v10 and 37348-2022051311067405 v5) from the Ministry of Research. Their sanitary status was assayed monthly, the animals were free of known pathogens and parasites. Breeding couples of clean *Psammomys obesus* are available on demand.

### Guide RNA design

RNA guides for the CRISPR-Cas9 strategy were designed and optimization was performed in collaboration with personnel at the IGBMC/PHENOMIN-ICS. The *Psammomys obesus Abca4* gene sequence was aligned against *Rattus norvegicus* (identity 94%) and *Mus musculus* (identity 92%) *Abca4* gene sequences. Deletion of exon 5 was chosen to obtain a frame shift and gene deletion. The guide RNAs were searched for using the CRISPOR web site (http://crispor.tefor.net/crispor.py) according to the instructions. Two guide RNAs were selected, the 5’ gR58 guide and the 3’ gR80 guide. Both should cut close to the splice acceptor or donor sites of exon 5 in order to increase cut efficiency. The guides were synthesized and tested *in vitro* (on the PCR fragment containing the target sequence and in the presence of the Cas9 protein).

### Plasmid construction

The pAAV-CMVep-HA-SpCas9 plasmid was constructed by inserting by SLIC (Sequence and Ligation Independent Cloning) the NLS-SpCas9-NLS sequence, amplified from pLentiCRISPRV1 plasmid [38] with an HA tag at its N-terminal end, in the pAAV-MCS plasmid (Stratagene) downstream of the CMV promoter. In order to reduce the size of the pAAV, the hGHpolyA signal of the pAAV-MCS plasmid was replaced by a small synthetic polyA sequence. For the construction of the pAAV-U6-gR58-U6-gR80-CAG-eGFP, the two guide RNAs R58 (TTTCACTCATGTGCAGGAAG) and R80 (GGTAAGTGAGGCCTGTTAAG) selected as targeting both 5’ and 3’ of exon 5 (a critical exon), were separately cloned in the BpiI site of the intermediate plasmid pSL1190-U6-lacZ-trRNA (Molecular Biology and Virus Service, IGBMC, Illkirch, France). The two U6-gRNA-trRNA cassettes were amplified and cloned by assembly (NEBuilder, New England Biolabs) in a pAAV already containing a pCAG-eGFP cassette.

### AAV production

Recombinant adeno-associated virus (AAV) serotype 8 (AAV8) were generated by a triple transfection of HEK293T/17 cell line using Polyethylenimine (PEI) transfection reagent and the 3 following plasmids: pAAV-U6-gR58-U6-gR80-CAG-eGFP or pAAV-CMV-HA-spCas9, pAAV2/8 (Addgene 112864 deposited by Dr Wilson, Penn Vector Core) encoding the AAV serotype 8 capsid and pHelper (Agilent) encoding the AV helper functions. 48h after transfection rAAV8 vectors were harvested from cell lysate and treated with Benzonase (Merck) at 100U/mL. They were further purified by gradient ultra-centrifugation with Iodixanol (OptiprepTM density gradient medium) followed by dialysis and concentration against Dulbecco’s Phosphate Buffered Saline (DPBS) using centrifugal filters (Amicon Ultra-15 Centrifugal Filter Devices 100K, Millipore). Viral titres were quantified by Real-Time PCR using the LightCycler480 SYBR Green I Master (Roche) and primers targeting eGFP (5’- GACGACGGCAACTACAAGA-3’, 5’-CATGATATAGACGTTGTGGCT-3’) or spCas9 (5’- GCATCCTGCAGACAGTGAAG-3’, 5’-TTCTGGGTGGTCTGGTTCTC-3’). Titres are expressed as genome copies (GC) per ml.

### Subretinal injections

*Psammomys obesus* were anaesthetized by inhalation of gaseous anesthetic (Isoflurane 2%) administered constantly through a small animal mask connected to a pump. The eyes were anesthetized by local anesthetic (Tetracaine) before surgery. The eyeball was exposed by pulling down the skin. A small incision was made through the *ora serrata* on the temporal side using the tip of a sharp 33-gauge needle. An injection needle (Hamilton syringe, 33 gauge blunt end) was inserted into the eyeball through the incision until slight resistance was felt, and ∼2µl of AAV vectors per eye were carefully injected into the subretinal space.

### Electroretinography (ERG)

*Psammomys obesus* were dark-adapted overnight, and the experimental procedures were done under complete darkness using an infra-red camera, as reported previously [21]. Animals were anesthetized with a subcutaneous injection of ketamine (25ng/kg) and medetomidine (7ng/kg). The pupils were dilated using Atropine 1% and the cornea was anesthetized with a drop of 0.5% Tetracaine. The scotopic ERG responses were evoked to stepped flashes of white light of increasing intensity, 0.0003-10.0 cd.s/m2 (rod responses and mix rod-cone response). Following scotopic recordings and after 10 minutes of light-adaptation (30 cd.m-2 background light), photopic (cone-mediated) ERGs were evoked using flashes of white light of 0.01-10.0 cd.s/m2. A 6-10-15-30-Hz photopic flicker response was also obtained using 1.0 cd.s/m2 and background illumination.

### Non-invasive Ocular Imaging

*Psammomys obesus* were anaesthetized by inhalation of isoflurane (IsoFlo®, Zoetis, Malakoff, France) administered constantly at 4% through a small animal mask in a 50/50 mix of air and O_2_ at 0.8l/min. A single drop of tropicamide 0.5% (Mydriaticum ®, Théa Pharma, Clermont-Ferrand, France) was used for pupil dilation, and the ocular surface was lubricated with either artificial tears (Artelac®, Bausch & Lomb, Montpellier, France) for OCT, or with ophthalmic gel (Ocry-Gel®, TVM, Lempdes, France) for fundus imaging. Retinas were first imaged with an Envisu R2200 SD-OCT (Bioptigen-Leica, Durham, USA), then with a Micron III fundus / fluorescence camera (Phoenix Research Laboratories, Pleasanton, CA, USA), during the same anaesthesia. Short sequences of fundus images were aligned, averaged and sharpened in Fiji [39, 40] using a macro developed by Mark Krebs (the Jackson Laboratory, Bar Harbor, MA, USA).

### mRNA quantification

For RT-PCR analysis, retinal RNA was isolated with RNeasy mini kit (Qiagen), following the manufacturer’s instructions including DNAse treatment. We modified slightly the first step, as dissociation of tissues was performed using a pestle with 500 μL of TriReagent solution. 100µl chloroform was directly added, and the preparation was transferred to a phase-lock gel tube after 15min incubation at room temperature. The gel tube was centrifuged 15 min at 12,000g and 4°C. To recover RNA, the supernatant was added to an equal volume of 70% ethanol. RT-PCR was performed from 700 ng of RNA with the high capacity RNA-to-cDNA kit. A second negative control was added, consisting of water, enzyme and RT buffer only, to confirm the absence of contamination. QPCR was performed in 96-well plates (Applied Biosystems), following the manufacturer’s instructions. Experiments were carried out on a 7300 Real-Time PCR System (Applied Biosystems) with a first step of denaturation for 5 min at 95 °C, followed by a cycle of 40 repetitions of 15s at 95°C, then 1 min at 60°C. After estimation of the sample concentration to use (between 1/10 and 1/100) and validation of primer efficiency (>80%), each sample was processed in duplicate and using a dilution range, for all studied genes: Abca4, rod-specific (Rho), cone-specific (Opn1mw), and housekeeping genes (Gapdh,Rplp0). See Supplementary Table 1 for primer references. Analysis was performed using the ΔΔCt method.

### Western Blotting

Protein from *Psammomys obesus* retina was extracted by sonicating in 100µl of lysis buffer (20 mM Tris-HCl pH7.6, 150 mM NaCl, 1% triton X-100, 1mM EDTA, 0.2% SDS) containing 1µl of proteinase inhibitor, as previously reported [21]. The lysate obtained was centrifuged 30 min at 13 000 rpm and 4°C, and the supernatant protein concentration was estimated by the Bradford method. Samples (20 μg protein/lane) were loaded and separated on SDS-PAGE gels, 90 min at 110V, following by semi-dry transfer to nitrocellulose membranes, 30min at 25V. After protein transfer, membranes were incubated in blocking solution (20mM Tris-HCl pH 7.6, 150mM NaCl, 0.1% Tween 20, 3% Bovine Serum Albumin), 1h at room temperature before overnight incubation at 4°C with the primary antibodies. After extensive washing in buffer, 4 x 5 min, membranes were incubated with secondary antibodies (rabbit anti-mouse IgG-HRP for monoclonal primaries, goat anti-rabbit IgG-HRP for polyclonal primaries), diluted 1:10,000 for 30min at room temperature. After washing again, 4 x 5 min, membranes were developed in the dark with the ECL Western Blotting Detection kit (Pierce), following the manufacturer’s instructions. Exposure times for each blot ranged between 5 to 30 seconds to obtain linear densitometric readings.

### Immunohistochemical staining

*Psammomys obesus* eyes were collected after euthanasia. Eyes were cut along the ora serrata to remove the cornea and lens and fixed 2 h at room temperature in 4% paraformaldehyde (PFA) in sucrose/phosphate-buffered saline (PBS). After extensive washing in PBS, dissected eyeballs were cryoprotected overnight and then embedded in OCT and sectioned by cryostat (10-12μm). *Psammomys obesus* retinal sections were permeabilized with 0.1% Triton x-100 for 5 min at room temperature, then incubated for 1h in blocking buffer (3% Bovine Serum Albumin, 0.1% Tween20 in PBS) at room temperature. As published previously [21, 22], primary antibodies were diluted in blocking buffer and incubated overnight at 4°C. Sections were washed three times in PBS and incubated with fluorochrome-conjugated secondary antibodies (1:1000) for 2h at room temperature. Secondary antibodies were donkey anti-rabbit or anti-mouse IgG conjugated with Alexa Fluor™ 488 or 647.Sections were washed again and then incubated 2 min with 1 4,6- diamidino-2-phenylindole (DAPI) (1:500). Sections were washed again and mounted with PBS-Glycerol 1:1. Images were captured using an inverted microscope (Zeiss Observer 7) equipped with objectives (63x, N.A. 1.4), a module for optical sectioning by structured illumination (Zeiss ApoTome.2) and a digital camera (Hamamatsu ORCA-Flash 4.0).

To prepare flatmounts of retina, *Psammomys obesus* eyes were enucleated as quickly as possible after euthanasia and immediately placed in 4% paraformaldehyde (PFA) as above. Retinas were gently detached from the eyecup, washed with PBS and mounted in Mowiol® 4-88, with the photoreceptor layer facing up. Imaging was conducted on a NanoZoomer S60 Digital slide scanner.

### Transmission electron microscopy

*Psammomys obesus* eyes were fixed in 2.5% glutaraldehyde in 0.1M sodium cacodylate buffer for 24h-48h. Samples were washed with 0.1M sodium cacodylate buffer and post-fixed with osmium tetroxide 0.5% in water, then dehydrated in an ethanol series (25%-50%-70%- 90%-100%2X) (15 min each) and propylene oxide 2x15 min and embedded in Epon EMbed 812 resin. For electron microscopy, ultrathin sections (70-80nm) were cut with an Ultracut EM FC7 Leica. Sections were collected on 200 mesh copper grids and stained in 1% uranyl acetate and observed using a Hitachi H 7500 transmission microscope equipped with an AMT Hamamatsu camera (Tokyo Japan).

### UPLC-MS analysis of bisretinoids

*Psammomys obesus* eyes derived from either control injected or Crispr-Cas9 injected animals (n=4 each group) were sent for analysis by ultra-high performance liquid chromatography (UPLC) coupled with mass spectrometry (MS). The person performing the analysis was blinded to donor origin. Using previously published methods [41], compound elution was achieved using an Alliance HPLC (Waters Corp Milford, MA, USA) with a Delta Pak[circlecopyrt]R C4 (5 μm, 3.9 × 150 mm; Waters) or an AtlantisR dC18 (3 μm, 4.6 × 150 mm; Waters) column. The mobile phase for the C4 column a gradient of acetonitrile in water with 0.1% trifluoroacetic acid: 0-5 min; 75% acetonitrile, flow rate, 0.8 ml min−1; 5-30 min, 75-100% acetonitrile; flow rate, 0.8 ml min−1; 30-35 min,100% acetonitrile, flow rate, 0.8-1.2 ml min−1; 35-50 min, 100% acetonitrile, flow rate, 1.2ml min−1. With the dC18 column a gradient of acetonitrile in water with 0.1% trifluoroacetic acid was utilized: 75-90% acetonitrile (0-30 min); 90-100% acetonitrile (30-40 min); 100% acetonitrile (40-100 min) with a flow rate of 0.5 ml min−1. Absorbance (Waters 2996 Photodiode Array) and fluorescence (Water 2475 Multi λ Fluorescence Detector; 18 nm bandwidth) were detected at the indicated wavelengths. Fluorescence efficiency was estimated as fluorescence peak area (μV s)/absorbance peak area (μV s).

## Acknowledgements

The authors wish to thank Drs. M.P. Felder-Schmittbuhl and F. Pfrieger for constructive discussions during manuscript preparation; Ms. C. Rouyer for expert technical assistance for transmission electron microscopy; Dr. C. Sandu for valuable help with qPCR analyses; and Ms. N. ElKorti for immunochemical staining. The authors extend their deep gratitude to the funding agencies that made these experiments possible: UNADEV/ITMO Aviesan 2018- 2021, Fondation de France (Association Berthe Fouassier) and USIAS (DH); (JRS).

