## supplementary figures and table for "Abca4 inhibition in a cone-rich rodent leads to Stargardt Disease type 1-like retinal degeneration"

Supplementary Figure Legends

Figure 1. Cas9 and eGFP expression in *Psammomys obesus* following subretinal injection of rAAV-CRISPR/Cas9. (**A-B**) Representative retinal whole-mounts of sgRNA/eGFP control (A) and sgRNA/CRISPR/Cas9 (B). A control animal 7d post-injection (DAPI labelling A') shows eGFP expression specifically in PRs (A'') but with no Cas9 expression (A'''). Sections of a treated animal 7d post-injection (DAPI labelling B') again shows eGFP expression specifically in the PR layer (B''), indicating transduction by the rAAV-sgRNA vector. There is also distinct staining for Cas9 in PRs present in the ONL (B'''), indicating successful transduction by both rAAV. Abbreviations: DAPI, 4′,6-diamidino-2-phenylindole; INL, inner nuclear layer; ONL, outer nuclear layer. Scale bar in B = 200µm for panels A and B, scale bar in B''' = 100µm for all other panels.

Figure 2. Representative scotopic single flash ERG recordings from individual *Psammomys obesus*. Shown on the left, ERGs were recorded across a range of increasing light flashes (0.03-$10^4$ mcd/s/m2) under scotopic (dark adapted) conditions with sgRNA/eGFP-injected right eye (control) compared to the sgRNA/CRISPR/Cas9-injected left eye (test). A, low intensity flashes (traces 1-3-5-7 = control right eye, R; traces 2-4-6-8 = test left eye, L); B, medium intensity flashes (traces 1-3-5 = control right eye, R; traces 2-4-6 = test left eye, L); C, high intensity flashes (traces 1-3-5 = control right eye, R; traces 2-4-6 = test left eye, L). Shown on the right, numerical values for "a" and "b" wave amplitudes (µV) for each trace corresponding to different intensities (A', B', C'). Whereas no differences in these values are seen below ~0.1 mcd, the test eye shows systematically lower values as flash intensity increases. Y-axis on ERG panels, divisions = 100µV; x-axis on ERG panels, divisions = 50 msec. The blue arrow shows appearance of the "i" wave, thought to represent ganglion cell activity and characteristic of *Psammomys* and human ERG traces.

Figure 3. Representative photopic single flash ERG from individual *Psammomys obesus.* ERGs were recorded across a range of light intensities under photopic conditions with sgRNA/eGFP control right eye (A and B, left) compared to the sgRNA/EGFP/Cas9 test left eye (A and B, right). Lower amplitude responses were obtained with increasing intensities of flash stimuli in the treated left eye compared to the control right eye. Numerical values for "a" and "b" wave amplitudes (µV) for each trace corresponding to different intensities (A’, B’). The test eye shows systematically lower values as flash intensity increases. Y-axis on ERG panels, divisions = 100µV; x-axis on ERG panels, divisions = 25 msec. The blue arrow shows appearance of the “i” wave, thought to represent ganglion cell activity and characteristic of *Psammomys* and human ERG traces.

Figure 4. Representative photopic flicker ERG from individual *Psammomys obesus.* ERGs were recorded across a range of frequencies (5-30 Hz) and at 10 cd/s/m$^2$ under photopic conditions, with sgRNA/eGFP control right eye (left) compared to the sgRNA/CRISPR/Cas9 test left eye (right). Lower amplitude responses were seen at all flicker frequencies in the treated left eye compared to the control right eye. Beneath the traces are shown numerical values for flicker response amplitudes (difference between trough N1 and peak P1, given in µV) for each trace. The test eye shows systematically lower values. Y-axis on ERG panels, divisions = 100µV; x-axis on ERG panels, divisions = 20 msec.

Figure 5. Mean ERG amplitudes recorded across a range of light flashes under scotopic (A, B), photopic (C, D) and photopic flicker (E) conditions from rAAV-sgRNA/eGFP control compared to unoperated control *Psammomys obesus* (60d post-injection). There were no significant differences between the two groups for any condition: A, scotopic “a” wave; B, scotopic “b” wave; C, photopic “a” wave; D, photopic “b” wave; E, photopic flicker. Open circles, rAAV-injected animals; closed circles, unoperated animals. Data represented as mean ± SEM. Statistical values between sgRNA/eGFP control eyes and unoperated eyes were calculated using two way ANOVA.

Figure 6. ERG recordings from *Psammomys obesus* at 90d and 210d after injection of rAAV-CRISPR/Cas9. Scotopic (A, B) and photopic (C, D) ERGs were recorded across a range of light flash intensities on sgRNA-CRISPR/Cas9-injected animals at 90 and 210 days post-injection. Mean scotopic “a” (A) and “b” waves (B) did not show any statistically significant differences between the two groups; on the other hand, under photopic conditions significantly lower amplitudes were seen in eyes after 210d compared to 90d (“a” wave, C; “b” wave, D). p values in italics comparing the maximum intensity flash at the two ages. Data represented as mean ± SEM. *$p < 0.05$; **$p < 0.01$. Significance values between 90 and 210d were calculated using two way ANOVA.

Figure 7. Fundus imaging of sgRNAs-eGFP control and CRISPR/Cas9-treated eyes at 60d post-injection. Control eyes showed a small lesion corresponding to the injection site (white arrow in A, C), but otherwise showed smooth uniform bluish retina with blood vessels emanating from the optic nerve head (ONH); B, D, fundus imaging of CRISPR/Cas9-treated eyes showing extensive loss of retina with pigmented patches and visible blood vessels.

Figure 8. Fundus imaging of CRISPR/Cas9-treated and unoperated eyes at 210 days post-injection. A: Fundus imaging of control eyes showed smooth uniform bluish retina with blood vessels emanating from the optic nerve head (ONH); A': fluorescence angiography of control eyes showed normal vasculature with retinal arteries and veins emerging from the ONH. B: Fundus imaging of CRISPR-Cas9-treated eyes showed extensive loss of retina with pigmented patches and visible blood vessels. B' Fluorescein angiography of CRISPR-Cas9-treated eye shows degenerated retina and abnormal vessels. B'', B''': Optical Coherence Tomography (OCT) imaging of CRISPR-Cas9-treated retinas showed a complete disappearance of retina with, on the edge of damage site, a complete disappearance of the ONL, while vestigial inner retina was still present.

Figure 9. Fundus imaging of sgRNAs/eEGFP control and CRISPR/Cas9-treated eyes at 30 and 60d post-injection. sgRNAs/eGFP control eyes were normal at 30 (A) and 60d (B). Fundus autofluorescence imaging of sgRNAs/eGFP control eyes showed stable eGFP expression at 30 (C) and 60d (D), without visible changes over time. OCT imaging of sgRNAs/eGFP control eyes showed the normal laminated retinal structure, with a brightly reflective retinal pigmented epithelium (RPE) and dark cell layers (INL, inner nuclear layer; and ONL, outer nuclear layer) with the GCL (ganglion cell layer) separated by a wide plexiform layer at 30 (E, G) and 60d (F, H), without changes over time. Fundus imaging of CRISPR/Cas9-treated eyes showed the presence of a lesion corresponding to the injection site at 30 (I) and 60d (J), with an enlargement of the damage site at 60d (delimited by dashed line). Fundus autofluorescence imaging of CRISPR/Cas9-treated eyes showed broad eGFP expression at 30 (K) and 60d (L), with an enlargement of the hypofluorescent damage site at 60d (delimited by dashed line). OCT imaging of CRISPR/Cas9-treated eyes showed degeneration of retina tissue with enlargement of the lesion between 30 (M) and 60d (N), delimited by dashed line. Transverse OCT sections made at three different levels across the lesion site showed that disappearance of the ONL progressed between 30 (O) and 60d (P), indicated by length of white bars beneath lesions.

Supplementary Table Legends

Table 1: PCR primers used for verifying Abca4 knockdown and quantification.

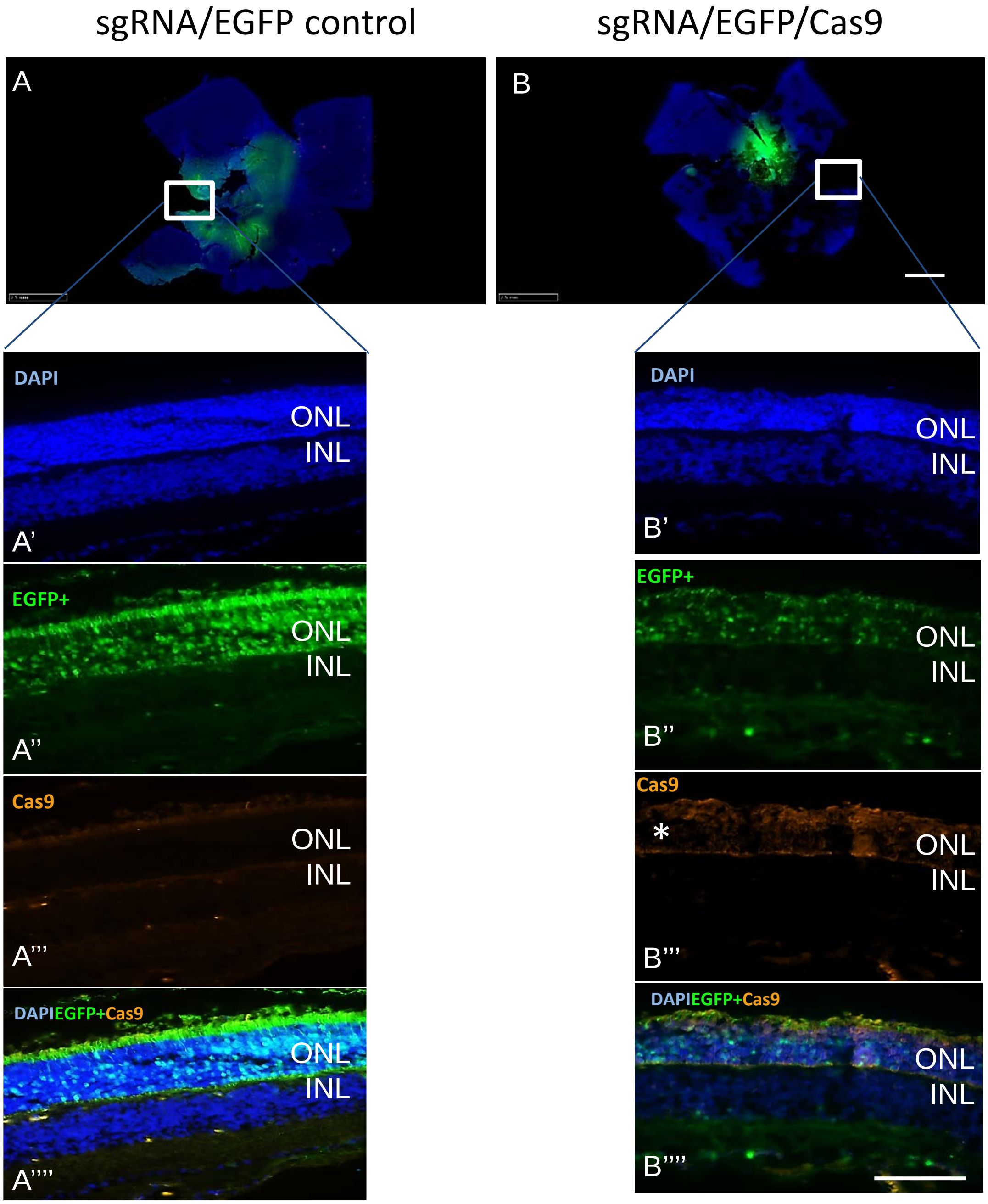
sgRNA/EGFP control
sgRNA/EGFP/Cas9
A
B
DAPI
ONL
INL
A'
EGFP+
A''
Cas9
A'''
DAPIEGFP+Cas9
A''''
B'
B''
*
B'''
B''''

### Scotopic single flash ERG

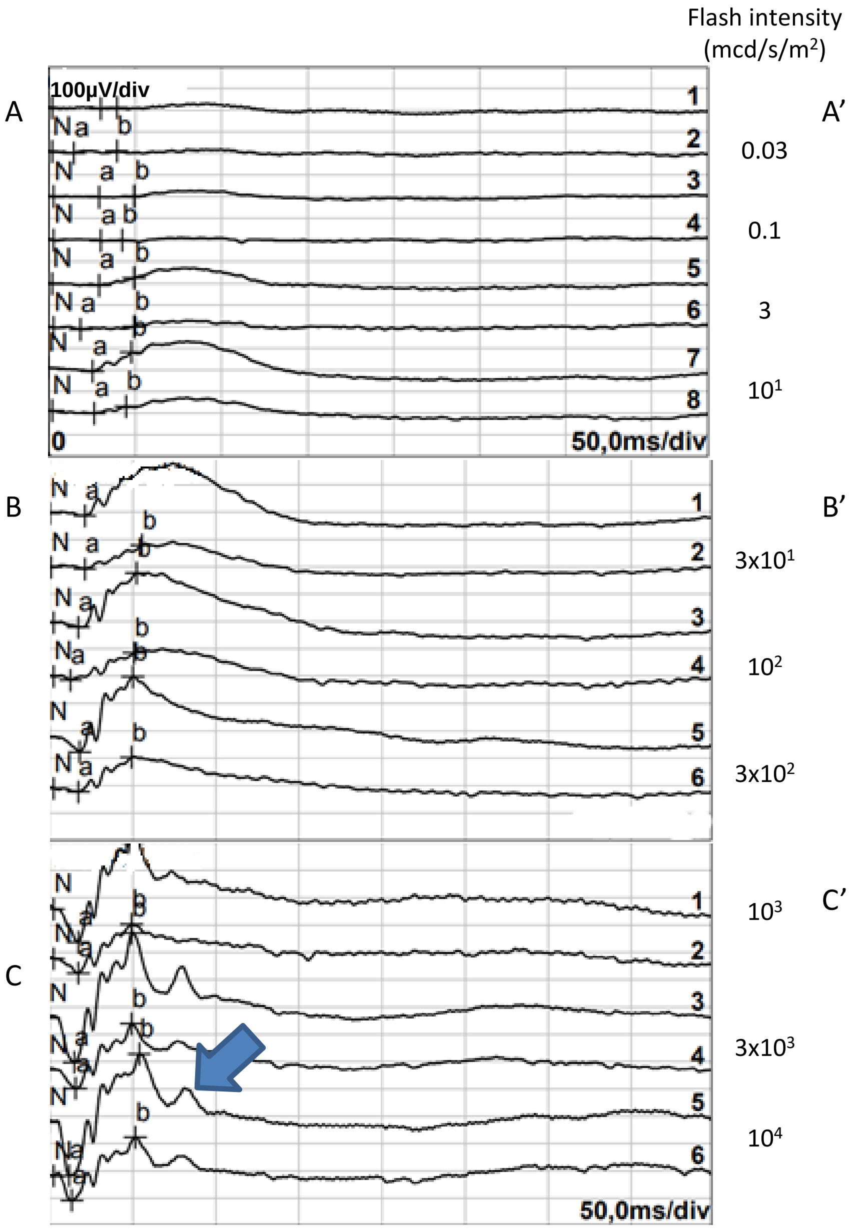


Amplitudes (µV)

A'

| Channel | a-wave | b-wave |
|---|---|---|
| 1 R-1 | 4,00µV | 4,66µV |
| 2 L-2 | 6,40µV | 8,79µV |
| 3 R-1 | 2,76µV | 3,86µV |
| 4 L-2 | 2,47µV | 1,22µV |
| 5 R-1 | 7,25µV | 35,4µV |
| 6 L-2 | 8,59µV | 13,4µV |
| 7 R-1 | 15,4µV | 88,7µV |
| 8 L-2 | 12,1µV | 31,3µV |

B'

| Channel | a-wave | b-wave |
|---|---|---|
| 1 R-1 | 9,33µV | 177µV |
| 2 L-2 | 7,28µV | 87,4µV |
| 3 R-1 | 16,4µV | 198µV |
| 4 L-2 | 13,0µV | 99,7µV |
| 5 R-1 | 53,5µV | 274µV |
| 6 L-2 | 12,5µV | 126µV |

C'

| Channel | a-wave | b-wave |
|---|---|---|
| 1 R-1 | 121µV | 376µV |
| 2 L-2 | 53,3µV | 179µV |
| 3 R-1 | 161µV | 471µV |
| 4 L-2 | 67,6µV | 237µV |
| 5 R-1 | 196µV | 445µV |
| 6 L-2 | 89,7µV | 231µV |

Photopic single flash ERG

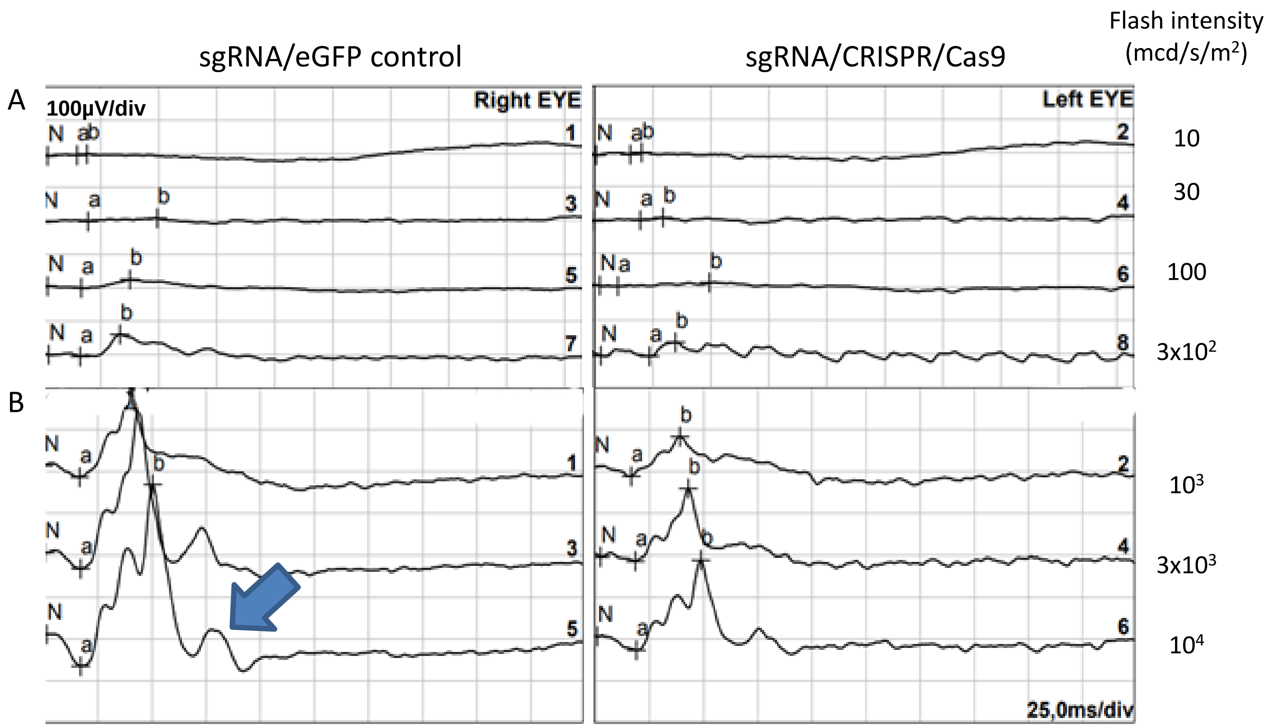


A'

| Channel | a-wave | b-wave |
|---|---|---|
| 1 R-1 | 1,76µV | 2,73µV |
| 3 R-1 | 1,83µV | 9,67µV |
| 5 R-1 | 4,66µV | 24,9µV |
| 7 R-1 | 5,18µV | 66,1µV |
| 2 L-2 | 1,64µV | 6,42µV |
| 4 L-2 | 1,86µV | 8,54µV |
| 6 L-2 | 1,66µV | 10,4µV |
| 8 L-2 | 2,37µV | 42,6µV |

B'

| Channel | a-wave | b-wave |
|---|---|---|
| 1 R-1 | 27,6µV | 203µV |
| 3 R-1 | 37,7µV | 382µV |
| 5 R-1 | 76,3µV | 434µV |
| 2 L-2 | 23,4µV | 95,5µV |
| 4 L-2 | 10,1µV | 173µV |
| 6 L-2 | 33,9µV | 216µV |

Photopic flicker ERG

sgRNA/eGFP control | sgRNA/eGFP/Cas9 | Frequency (Hz)

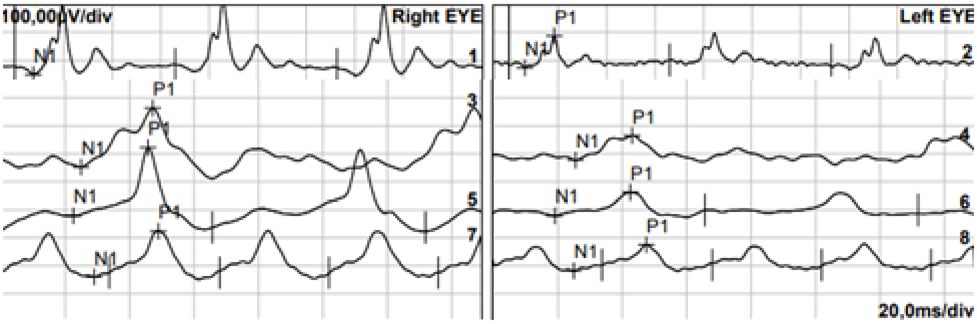


5

10

15

30

| Channel | N1-P1 |
|---|---|
| 1 R-1 | 248µV |
| 2 L-2 | 113µV |
| 3 R-1 | 211µV |
| 5 R-1 | 243µV |
| 7 R-1 | 165µV |
| 4 L-2 | 88,0µV |
| 6 L-2 | 83,1µV |
| 8 L-2 | 94,0µV |

ERGs from injected control AAV (sgRNA/eGFP only) vs unoperated control *Psammomys obesus*

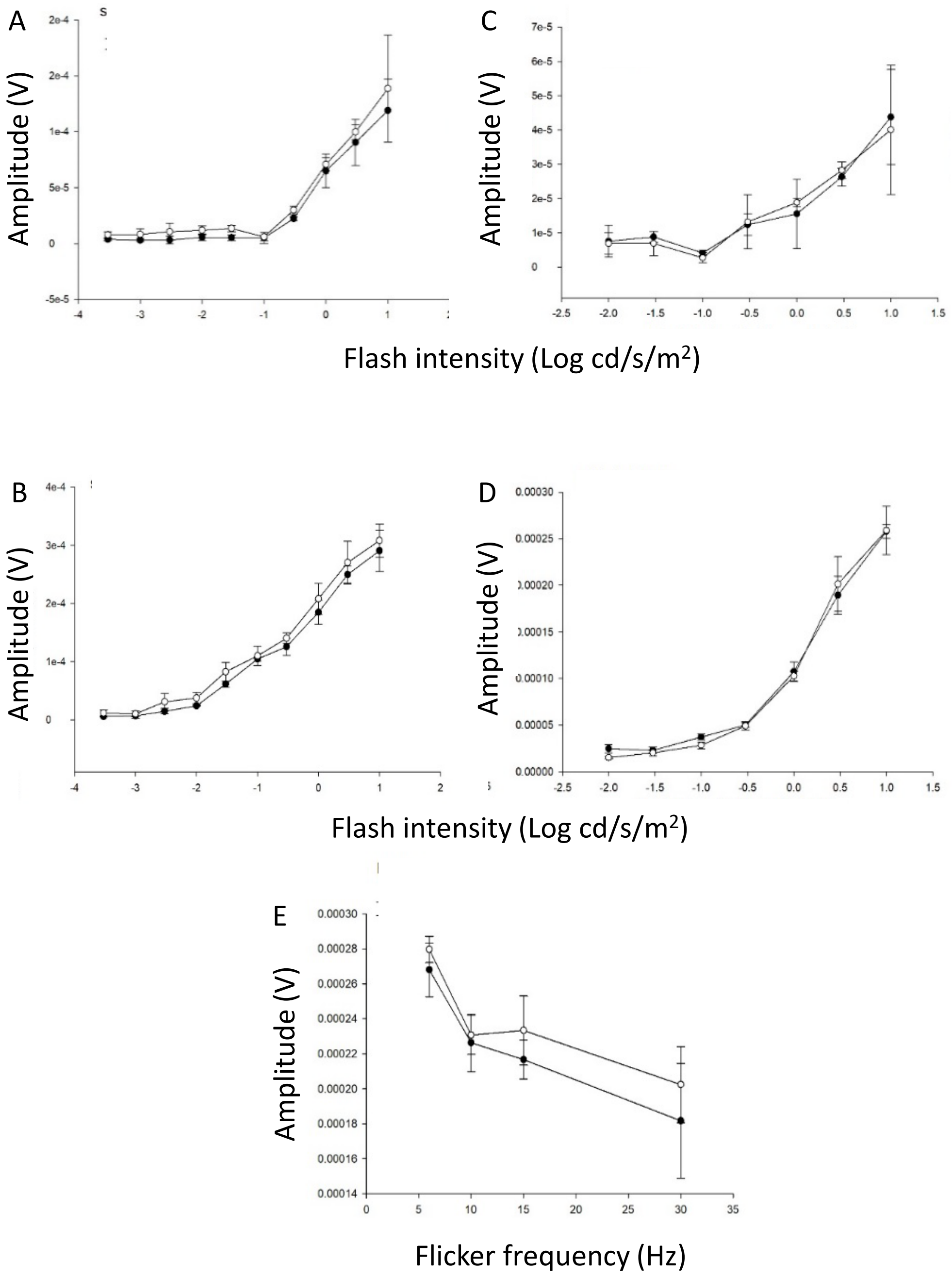

Comparison of ERG recordings made from *Psammomys obesus* at 90 and 210d after injection of rAAV-CRISPR/Cas9

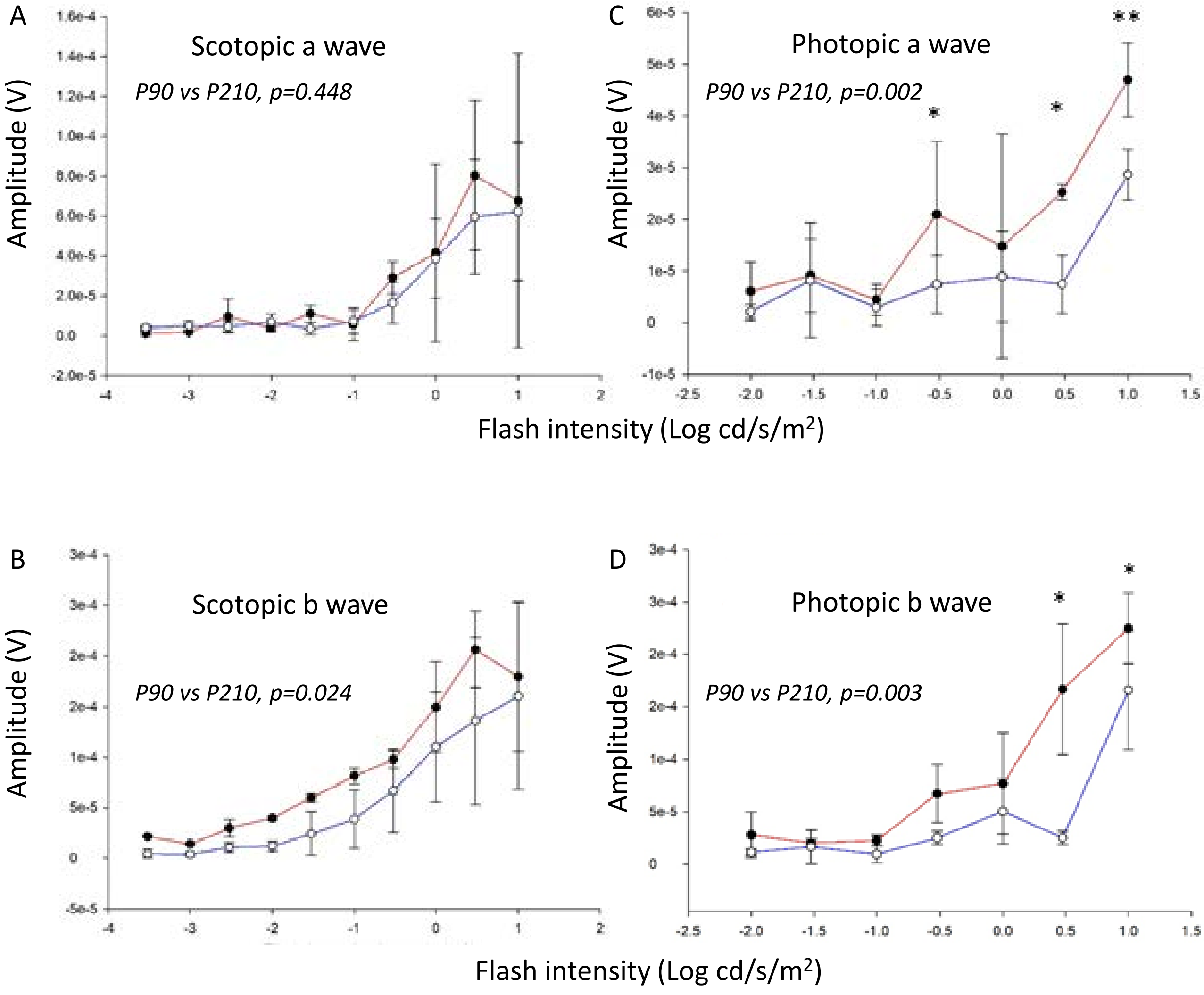

*Psammomys obesus*, 60 days post-injection

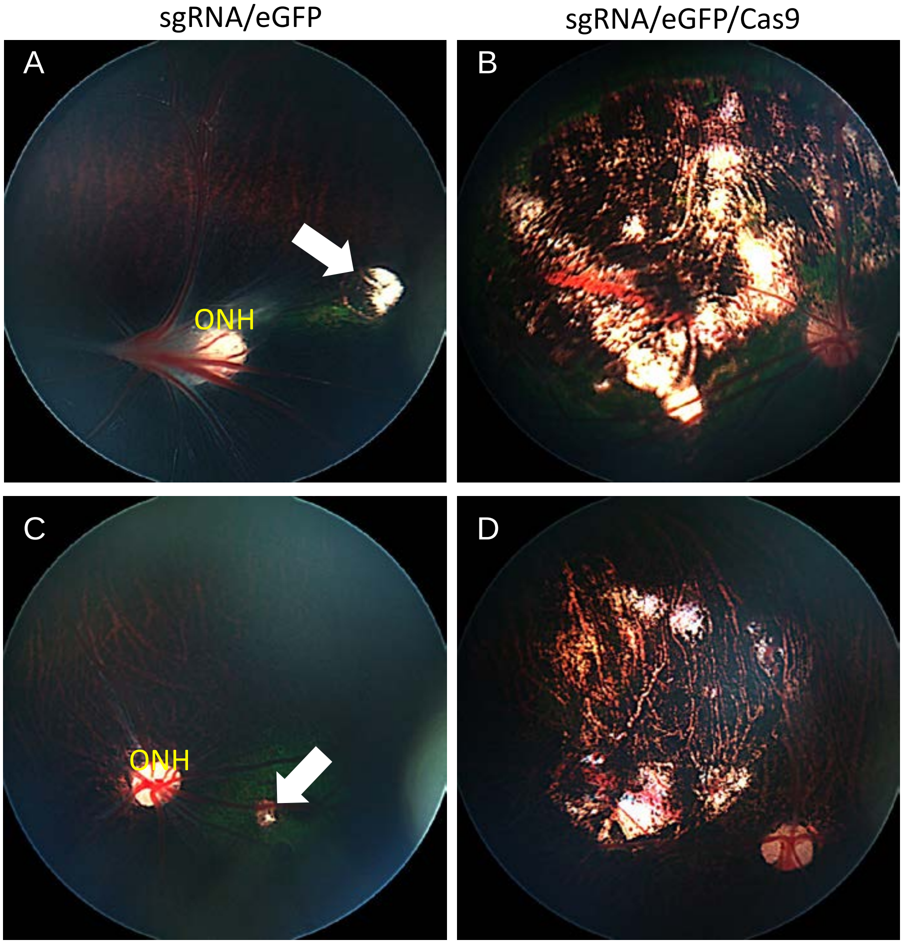

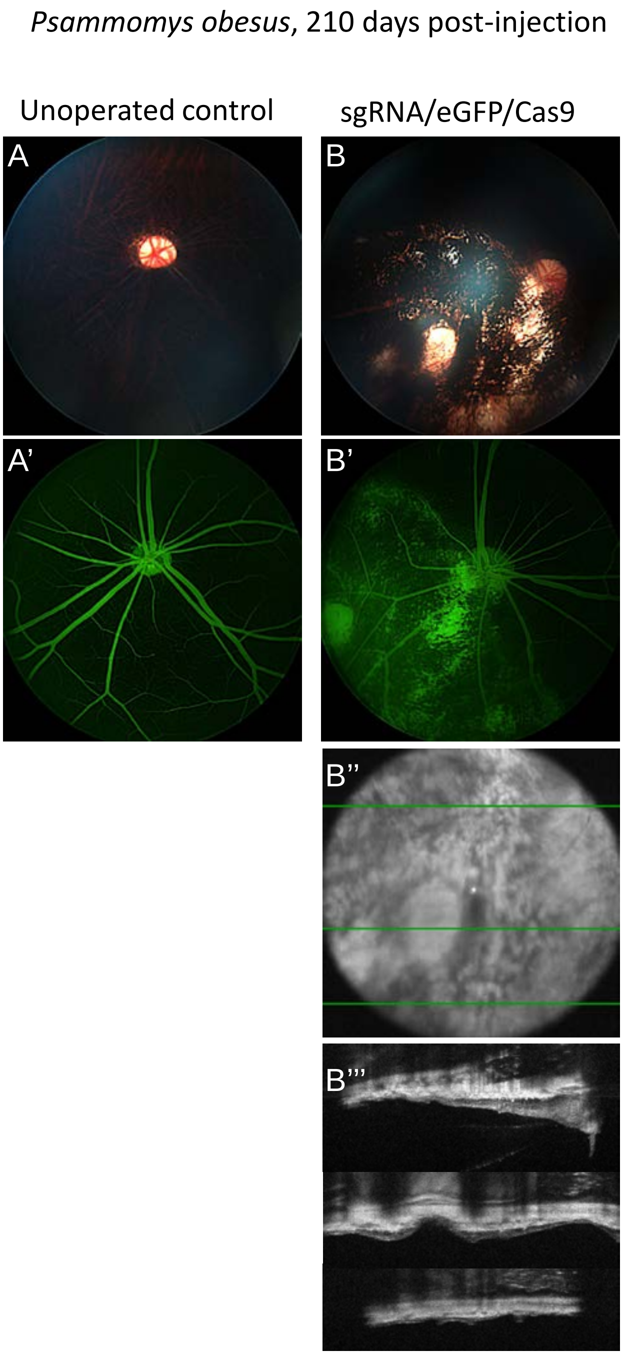
*Psammomys obesus*, 210 days post-injection
Unoperated control
sgRNA/eGFP/Cas9
A
B
A'
B'
B''
B'''

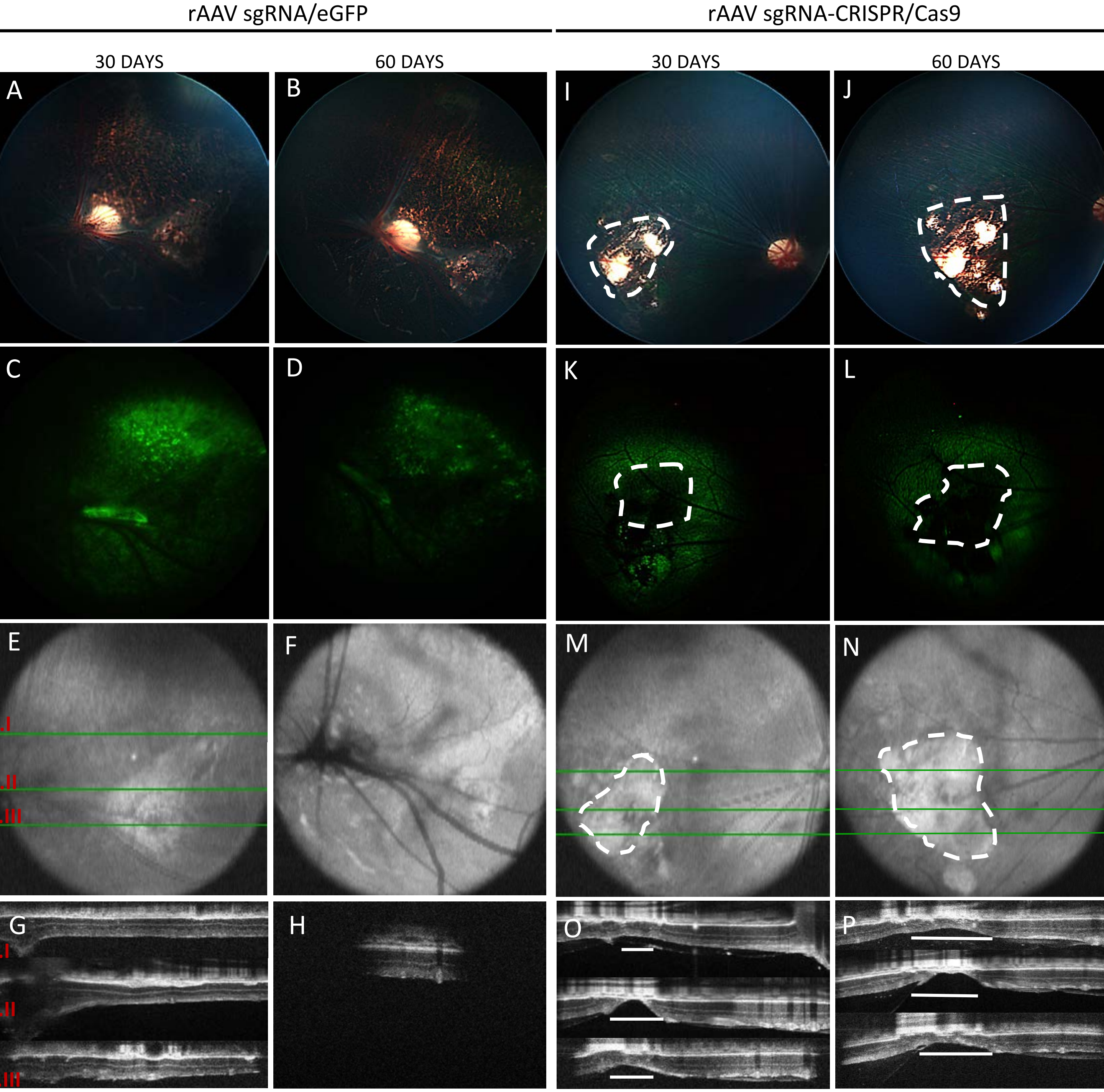

rAAV sgRNA/eGFP
rAAV sgRNA-CRISPR/Cas9
30 DAYS
60 DAYS
30 DAYS
60 DAYS
A
B
I
J
C
D
K
L
E
F
M
N
.I
.II
.III
G
H
O
P
.I
.II
.III

Table 1: Primers used to amplify *Psammomys obesus* sequences.

| | |
|---|---|
| *Psammomys* primers for gRNA verification | |
| Abca4 *Psammomys* F | CTTCTGTAGTTCCTTACCCTTCAC |
| Abca4 *Psammomys* R | GGTAGGTTCAGCTTCACGATT |
| *Psammomys* primers for mRNA quantification | |
| Abca4 ex48-49 F | GGATCTTCCAGCTGCTCATT |
| Abca4 ex48-49 R | GGAGGTCATAGGTCTCAGTCT |
| RHO F | CCATCCCTTTGACCATCATCT |
| RHO R | ATGCGTGTGACTTCCTTCTC |
| Opn1mw F | TGGCAATATGAGATTTGATGCT |
| Opn1mw R | CCAGACCCAAGAAAAGATGAT |
